# In Silico Treatment: a computational framework for animal model selection and drug assessment

**DOI:** 10.1101/2024.06.17.599264

**Authors:** Sergio Picart-Armada, Kolja Becker, Marc Kaestle, Oliver Krenkel, Eric Simon, Stephan Tenbaum, Benjamin Strobel, Kerstin Geillinger-Kaestle, Katrin Fundel-Clemens, Damian Matera, Kathleen Lincoln, Jon Hill, Coralie Viollet, Ruediger Streicher, Matthew Thomas, Jan Nygaard Jensen, Christian Haslinger, Holger Klein, Markus Werner, Heinrich J. Huber, Andre Broermann, Francesc Fernandez-Albert

## Abstract

The translation of findings from animal models to human disease is a fundamental part in the field of drug development. However, only a small proportion of promising preclinical results in animals translate to human pathophysiology. This underscores the necessity for novel data analysis strategies to accurately evaluate the most suitable animal model for a specific purpose, ensuring cross-species translatability. To address this need, we present *In Silico* Treatment (IST), a computational method to assess translation of disease-related molecular expression patterns between animal models and humans. By simulating changes observed in animals onto humans, IST provides a holistic picture of how well animal models recapitulate key aspects of human disease, or how treatments transform pathogenic expression patterns to healthy ones. Furthermore, IST highlights particular genes that influence molecular features of pathogenesis or drug mode of action. We demonstrate the potential of IST with three applications using bulk transcriptomics data. First, we assessed two mouse models for idiopathic pulmonary fibrosis (IPF): one involving injury with intra-tubular Bleomycin exposure, and the other Adeno-associated-virus-induced, TGFβ1-mediated tissue transformation (AAV6.2-TGFβ1). Both models exhibited gene expression patterns resembling extracellular matrix derangement in human IPF, whereas differences in VEGF-driven vascularization were observed. Second, we confirmed known features of non-alcoholic steatohepatitis (NASH) mouse models, including choline-deficient, l-amino acid-defined diet (CDAA), carbon tetrachloride hepatotoxicity injury (CCl_4_) and bile duct ligation surgery (BDL). Overall, the three mouse models recapitulated expression changes related to fibrosis in human NASH, whereas model-specific differences were found in lipid metabolism, inflammation, and apoptosis. Third, we reproduced the strong anti-fibrotic signature and induction of the PPARα signaling observed in the Elafibranor experimental treatment for NASH in the CDAA model. We validated the contribution of known disease-related genes to the findings made with IST in the IPF and NASH applications. The complete data integration IST framework, including an interactive app to integrate and compare datasets, is made available as an open-source R package.

**Author summary:** Preclinical testing plays a pivotal role in the drug development process, serving as a crucial evaluation phase before a new drug can be tested on humans in clinical trials. The drug must undergo a rigorous evaluation in *in vivo* and *in vitro* preclinical studies to assess its safety and efficacy. However, positive outcomes in preclinical animal models do not always translate to positive results in humans, mainly due to biological differences. Therefore, selecting an animal model that closely mirrors human disease traits and detecting and accounting for model limitations is of paramount importance.

Over the last decade, the availability of gene expression data in both animals and humans has substantially increased. Gene expression states and perturbations are routinely employed as a proxy to predict and understand changes in disease states. Here, we developed In Silico Treatment, a computational method designed to overlay the gene expression changes observed in animals onto humans, quantifying the change in human disease status. We applied this method to mouse models for idiopathic pulmonary fibrosis and non-alcoholic steatohepatitis, two severe fibrotic diseases. We successfully identified known features of the disease models and provide a granular gene-level rationale behind our predictions. Consequently, our method shows promise as an effective approach to improve animal model selection and thus clinical translation.

## Introduction

Animal models play a crucial role in improving understanding of human disease. Accordingly, drug development often relies on successful animal studies before proceeding to costly and lengthy clinical trials (Mak, Evaniew, and Ghert 2013). However, not all potential therapeutic concepts successfully translate from rodent and other animal models to humans, implying significant differences in molecular mechanisms across species that drive pathophysiology (McGonigle and Ruggeri 2014). As a result, the choice of the most appropriate animal model to study specific molecular and systemic modes of action is not straightforward, but requires a trade-off between ethical aspects regarding animal experimentation, financial and feasibility considerations, and animal model suitability to mimic the human disease (Breschi, Gingeras, and Guigó 2017; Wendler and Wehling 2010).

Important for the choice of suitable animal models is to understand if and how key mechanisms of pathology translate between species (Perel et al. 2007). While a given animal model may faithfully capture certain aspects of human disease, other disease-relevant mechanisms may be only poorly resembled and may require interrogation of a different model. In this regard, the quantification of model suitability from molecular readouts remains an open issue. For example, past studies have led to conflicting conclusions of low (Seok et al. 2013) or high resemblance (Takao and Miyakawa 2015) between murine models and human inflammatory diseases. Taken together, we believe there is a promising potential for *in silico* approaches to systematically gather knowledge on the aspects of a human disease that are well reflected in each specific animal model, facilitating a more targeted approach to increase the probability of success in subsequent experiments (Michelson and Reuter 2019). While attempts in this direction exist, so far there is no consensus on how to automate the assessment of animal model suitability on a molecular or transcriptome-wide level.

Here, we introduce *In Silico* Treatment (IST), a computational framework for the integrative analysis of human and *in vivo* animal model transcriptomics data. IST uses predictive modelling methods to quantify the overlap of ortholog gene expression changes between human patients and disease models for a particular human disease and molecular pathway. Besides comparing the suitability of specific animal models, IST also provides a framework to predict whether a particular drug treatment can potentially revert disease-related molecular profiles in humans. Furthermore, IST includes features supporting the interpretation of the gene signatures that reflect pathophysiology and treatment in disease models by helping evaluate them in the human context. Thereby, IST provides an integrative picture of human and disease model data at different levels including pathway (gene set) and gene-wise granularity.

We showcase capabilities and features in IST by applying it to two human diseases: Idiopathic Pulmonary Fibrosis (IPF), and Non-alcoholic Steatohepatitis (NASH). Despite the broad usage of animal models in IPF and NASH, the agreement and the resulting predictability between human and mouse gene expression changes is unknown, and thus the ability to draw conclusions from the molecular profiles remains elusive. In this context, we demonstrate how IST (i) determines which disease models for IPF and NASH most appropriately capture human gene expression changes on a pathway level helping select the most suitable animal model for pre-clinical research, (ii) evaluates potential treatments for a human disease by predicting the recovery of the healthy human molecular phenotype for each treatment on each pathway, and (iii) provides gene-level quantitative explanations behind the selection of a specific disease model or treatment compound.

## Results

### In Silico Treatment uses predictive modelling to compare the gene expression changes between *in vivo* models using a human disease reference

We used gene expression data in combination with the IST framework on IPF and NASH, two fibrotic human diseases, to compare a collection of frequently used *in vivo* mouse models for each of the indications and pathway of interest.

The IST data integration workflow requires the following input data: gene expression readouts from human control and disease samples, gene expression fold changes from each preclinical model, gene sets related to the human disease, and a gene orthology mapping that links the genes in the preclinical organisms to their human orthologs. After the data integration process in IST, two main outputs are generated. Firstly, for every gene set, there is a single quantitative measure that shows how well each preclinical model captures the changes observed in the human reference within the gene set. Secondly, for every gene set and gene, there is a quantitative measure that indicates how the changes in that particular gene in the preclinical model contribute to the overall similarity of the preclinical model to the changes in the human reference.

The IST workflow consists of three steps: First, predictive machine learning models, here partial least squares, are fit to human gene expression data to discriminate between the control group and patients with disease (left panel, Figure 1A). Second, significant gene expression fold changes of preclinical models are simulated onto the ortholog genes of the human reference samples. This results in simulated samples, whose expression profiles have undergone the same changes that were observed in preclinical models (middle panel, Figure 1A). In a third step, preclinical models are evaluated by predicting the response, also called disease score, of simulated samples based on the fitted predictive model. This quantifies whether the simulated changes have brought the simulated samples closer or further from human disease states (right panel, Figure 1A).

**Figure 1.**
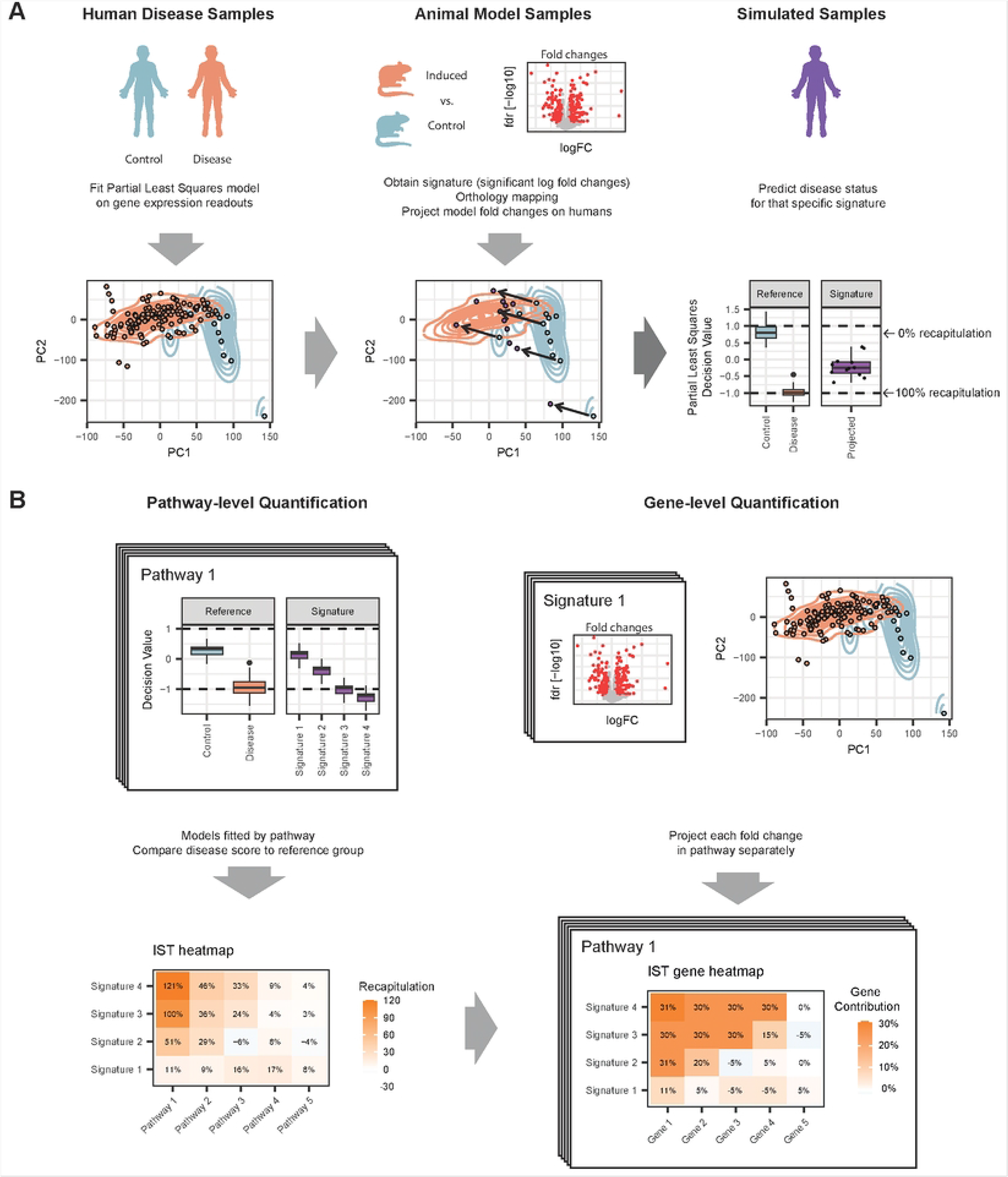
Overview of the In Silico Treatment workflow: (a) IST conceptual workflow. First, human disease samples are used to learn differences between healthy and disease gene expression patterns via predictive models. Second, fold changes of significantly deregulated genes in animal model signatures are overlaid onto the expression profile of their human ortholog genes, in the desired human population. This process is called the fold change simulation. Third, the newly obtained simulated human expression profiles are evaluated against the model from the first step. This resulting disease score is compared against disease scores of controls and disease. (b) Pathway models. Predictive models are fitted to gene sets representing key disease hallmarks. For each pathway and signature, the outcome of the IST workflow is expressed as percentage of ideal recapitulation. Signatures with recapitulations close to 0% describe a very modest modification of the disease score, while those closer to 100% indicate a switch of the expression profiles towards the desired human population. The pathway recapitulations can be decomposed into additive contributions per gene. IST thus identifies what genes in a signature positively and negatively contribute to the overall recapitulation, and how much.

Two alternative strategies to apply IST were devised, depending on whether to evaluate pathogenic effects in animal models or to predict the efficacy of disease treatment in humans. For the assessment of disease models, fold changes of gene expression from animal models relative to their respective controls are mapped onto human control samples. For the assessment of treatment, fold changes from treated animal models of disease relative to their untreated counterparts are mapped onto human disease samples. In both cases, a comparison of the predicted disease scores of simulated samples with that of human reference samples (disease or control samples, respectively) is performed. Disease scores are then expressed as the relative distance between simulated and human reference samples, with 100% representing ideal recapitulation and 0% no recapitulation at all (right panel, Figure 1A).

Regarding the outputs and graphical representations from the IST framework, it is possible to fit one disease score model for each gene set that represents a key disease pathway or feature. This enables IST to make granular choices for testing specific mechanisms or aspects of disease (left panel, Figure 1B). In addition, IST provides gene-level contributions by simulating each gene separately, to find agreeing and disagreeing gene expression patterns between disease model and human pathophysiology (right panel, Figure 1B). We provide an open-source implementation of the whole IST workflow using the R programming language.

### Comparison of the IPF disease models

IPF is a severe and fatal fibrotic lung disease of unknown cause, leading to aberrant lung tissue remodeling, excessive scarring, loss of tissue compliance and respiratory failure (Mari, Jones, and Richeldi 2019). Here, we used a reference IPF human dataset consisting of microarray gene expression readouts of lungs from control and IPF patients (Y. Wang et al. 2017). We then identified highly deregulated pathways in IPF by performing a gene set enrichment analysis (GSEA) (Subramanian et al. 2005) on the human reference data. We selected six disease-relevant pathways (Figure 2A), combining GSEA output and known disease pathomechanisms.

**Figure 2.**
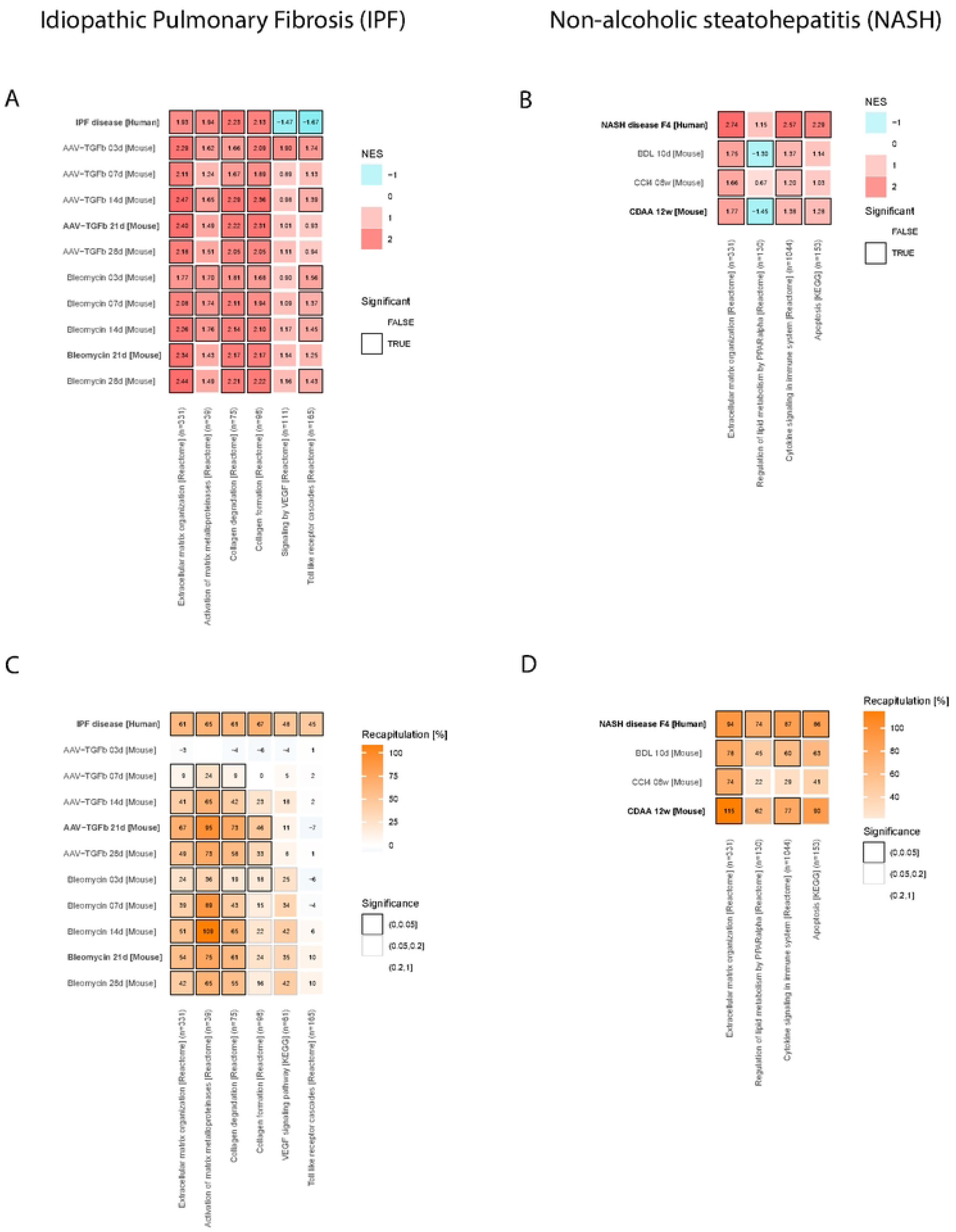
Assessment of animal model signatures for hallmark pathways in human IPF and NASH using GSEA and the IST pathway heatmap: **(A)** Gene set enrichment analysis (GSEA) of the human IPF signature and the animal model signatures, mapped to their ortholog human genes. The heatmap depicts normalized enrichment scores (NES) from a pre-ranked GSEA for six IPF-related pathways. The NES sign defines the direction of the enrichment (positive for upregulation, negative for downregulation). Significance for pathway deregulation indicated at 10% false discovery rate. **(B)** Analogously, pre-ranked GSEA of the human NASH signature and the animal model signatures. **(C)** IST pathway heatmap for IPF human data and animal models. Recapitulation percentages are displayed, being 0% no recapitulation (expression profiles after fold change simulation still look like healthy humans) and 100% ideal recapitulation (simulated expression profiles look like human IPF expression profiles). Significance for positive recapitulation indicated as false discovery rate ranges: from 0 to 5%, from 5% to 20%, and greater than 20%. **(D)** Analogously, IST pathway heatmap for NASH human data and animal models.

Multiple animal models for IPF have been established for pre-clinical research. Here we considered the models of intra-tracheal administration of mice with the cytostatic toxin Bleomycin, and a transgenic mouse model based on AAV6.2-induced overexpression of Transforming growth factor beta 1, or TGFβ1 (Strobel et al. 2015). Both the AAV-TGFβ1 and the Bleomycin mouse models were recorded across timepoints: 3, 7, 14, 21 and 28 days. After RNA sequencing, fold changes and significance were computed by timepoint. We also applied GSEA to the murine fold changes, mapping murine genes to their human orthologs (Figure 2A).

When applying the IST workflow across all selected pathways, the output for the IPF models showed low recapitulations within early expression changes in AAV-TGFβ1 mice (3d, 7d), not entailing sufficient molecular changes to resemble the human IPF gene expression data (Figure 2C). Conversely, later time points of the AAV-TGFβ1 model (14d, 21d and 28d) have larger resemblances to the human molecular signature, suggesting a delayed response in TGFβ1-mediated injury due to time required for viral transduction, conversion of the single-stranded AAV genome to transcriptionally active dsDNA, and actual gene expression. In alignment with this hypothesis and the corresponding lack of phenotypic changes (Strobel et al. 2022), we see only few differentially expressed genes at the 3d and 7d time points (Supplementary Figure 1C).

Aberrant extracellular remodeling, a key characteristic of several fibrotic diseases such as cardiac fibrosis, NASH, or IPF, is depicted in the extracellular matrix organization pathway in Figure 2C. IST demonstrated substantial agreement between human data with both intermediate and late time point AAV-TGFβ1 and all Bleomycin mouse model samples. The highest recapitulation of human data occurred at the 21d AAV-TGFβ1 model (67%) and the 21d Bleomycin mice (54%). For genes involved in the activation of matrix metalloprotease pathway, IST indicated large positive recapitulation values. Specifically, the highest recapitulation was observed in the AAV-TGFβ1 mouse model at 21d (95%), and the Bleomycin mouse model at 14d (109%), suggesting that these specific experimental conditions are most suitable for studying the activation of matrix metalloproteases in the context of lung fibrosis.

Important for extracellular matrix organization is a balance between collagen formation and collagen degradation. Interestingly, while the degradation of collagens was well represented by both IPF mouse models (Bleomycin 14d and AAV-TGFβ1 21d showing a recapitulation of 65% and 73% respectively), this was not the case for collagen formation where only AAV-TGFβ1 21d mice showed a sizeable recapitulation of 46%.

VEGF dependent tissue vascularization is an important factor in IPF pathology. VEGF signaling, originating mainly from airway epithelial cells, is typically moderate in the mature and heathy lung, while tissue damage and subsequent repair leads to re-vascularization (Barratt et al. 2018). Although targeting vascular endothelial growth factor (VEGF) has been approved as part of a triple kinase inhibition therapeutic strategy in IPF (Nintedanib, Boehringer Ingelheim, Germany), the role of VEGF signaling in IPF remains yet controversial. (Barratt et al. 2018)(Lee et al. 2008; Iyer et al. 2015)(Murray et al. 2017). While GSEA suggested pathway changes in opposite directions between disease models and human data (VEGF signaling pathway in Figure 2A), IST found a degree of agreement (Figure 2C), especially in the lung injury Bleomycin model (42% at 14d). Indeed, using animal model data from our facilities, when treating both mouse models with Nintedanib, lung vital capacity was only statistically significantly restored in the Bleomycin, but not in the AAV-TGFβ1 model (Supplementary Figure 1D), suggesting that the Nintedanib revertible phenotype in the prior mouse model better resembles the human pathology and its attenuation by Nintedanib.

Finally, we investigated innate immune signaling by toll-like receptor mediated pathways (pathway Toll-like receptor cascades, Figure 2C) which constitute important mediators of the inflammatory response in early tissue injury and remodeling (Karampitsakos et al. 2017). As a general picture, none of the mouse models show good resemblance of the human IPF data with respect to genes present in the TLR receptor pathway, with partially opposite changes in the 21d AAV-TGFβ1 model and the 3d and 7d Bleomycin model. This disagreement between animal models and human gene expression remains to be further investigated, begging the question whether additional disease models, apart from AAV-TGFβ1 or Bleomycin treated mice could be more suitable to study the effect of IPF on the innate immune system response.

### Comparison of NASH disease models

NASH, recently renamed to metabolic dysfunction-associated steatohepatitis (MASH), is a complication of non-alcoholic fatty liver disease (NAFLD) or metabolic dysfunction-associated steatotic liver disease (MASLD) (Rinella et al. 2023). NASH is an increasingly prevalent liver disease that can progress to cirrhosis and acute or chronic liver failure and is one of the most frequent indications for liver transplantation (Younossi et al. 2018). Hepatic steatosis due to long-term exposure of individuals to high fat and high-sugar diets is considered as one of the factors promoting NASH development. Within a fatty liver the associated liver cell damage and inflammation lead to progressively increasing fibrotic scarring caused by the excessive extracellular matrix deposition and finally cirrhosis and impaired liver function (Loomba, Friedman, and Shulman 2021). We used a human NASH reference with RNA sequencing data from liver tissue of individuals with increasing pathologically assessed fibrosis stages ranging from F0 to F4, i.e., from fatty liver with no fibrosis to marked fibrosis with cirrhosis (Pantano et al. 2021). We focused on assessing how murine models capture the molecular changes in F4 compared to F0. After running GSEA on this human data, and considering known disease pathomechanisms, we selected four pathways as examples for further examination (Figure 2A).

We considered three mouse models performed previously in our animal facilities complying with all necessary ethical and regulatory standards: the choline-deficient, l-amino acid-defined dietary model (CDAA) for 12 weeks, the carbon tetrachloride hepatotoxicity injury model (CCl_4_) for 8 weeks and the bile duct ligation (BDL) model at 10 days after surgery, which induces cholestasis and inflammation. Overall, these models are known to show different aspects of the pathology and varying degrees of clinical translatability (Hansen et al. 2017). Here, total mRNA was sequenced by standard NGS methods, fold changes were obtained for each animal model, and GSEA was applied after mapping murine genes to their human orthologs (Figure 2C).

Using IST, we studied key mechanisms of fibrosis progression in NASH through the gene set of extracellular matrix organization. All evaluated disease models aligned with human fibrosis stage 4 expression patterns (Figure 2B), especially CDAA (115%) followed by BDL (78%) and CCl_4_ (74%). These findings were expected since those three models are well described to study aspects of severe human liver fibrosis. Our focus on fibrosis stage 4 particularly fits with the CDAA choice, a sound model to study progression to NASH (Yanguas et al. 2016). Peroxisomes are subcellular organelles involved in β-oxidation of fatty acids as well as bile acid and cholesterol metabolism (Islinger, Cardoso, and Schrader 2010). Peroxisome proliferator-activated receptors (PPARs) are nuclear receptors regulating the proliferation of peroxisomes and consist of three subtypes, PPARα, PPARβ/δ and PPARγ. PPAR response genes are involved in glucose and lipid metabolism (Bougarne et al. 2018). IST suggests (Figure 2D) that lipid metabolism regulation by PPARα, as observed in NASH liver, was partially recapitulated in CDAA (62%), BDL (45%) and to a lesser extent in CCl_4_ (22%). The better recapitulation of lipid metabolism dysregulation in CDAA compared to CCl_4_ could be related to the chemotoxic fibrotic mode of action of CCl_4_, lacking certain metabolic aspects of NASH, as opposed to a diet-driven model like CDAA.

Inflammation during NASH progression is initiated by damaged liver cells and maintained by multiple immune cell types, such as tissue resident Kupffer cells as well as infiltrating immune cells. One key aspect is the release of inflammatory mediators, mainly cytokines and chemokines. In line, disease severity in NASH patients has been shown to correlate with the levels of inflammatory cytokines as IL1B, TNFα or IL6 (Plessis et al. 2016). Using IST, we found that cytokine immune signaling mechanisms are well recapitulated by common animal models of NASH (Figure 2D), especially in CDAA (77%) and BDL (60%) models. This aligns with known inflammatory features of the models: CDAA causes panlobular inflammation since week 3, and BDL’s bile acid accumulation promotes oxidative stress and necroinflammation (Yanguas et al. 2016).

The link between NASH and apoptotic pathways is well established. IST quantified the best recapitulation for CDAA (90%) and BDL (63%), followed by CCl_4_ (41%) (Figure 2D). IST thus distinguished signatures related to the type of cell death: the dietary nature of CDAA better aligned with cellular apoptosis as in human NASH, versus the injury by CCl_4_ administration, which induces necrosis rather than apoptosis (Manibusan, Odin, and Eastmond 2007).

### In Silico Treatment enables a gene-level evaluation of the disease model signatures

In the previous section, we used IST to compare different animal models in key disease pathways, aiming at optimal animal model selection. But the bare presence of sizeable differences between animal models within a disease pathway may not give sufficient granularity about the mechanistic reasons that could make one specific animal model more suitable.

In this section, we showcase the IST features that allow to compare different disease models by assessing the individual gene contributions behind the pathway recapitulation scores. For every signature, we quantified the contribution of each gene to the overall signature recapitulation by simulating each gene’s fold change onto humans separately. We will use these features to explain the rationale behind some of the recapitulation values that IST predicted for the IPF and NASH models. We discuss the fold changes of some key genes (Figures 3A and 3B) and how they translate into gene contributions (Figures 3C, 3D, 3E, 3F and 3G)

**Figure 3.**
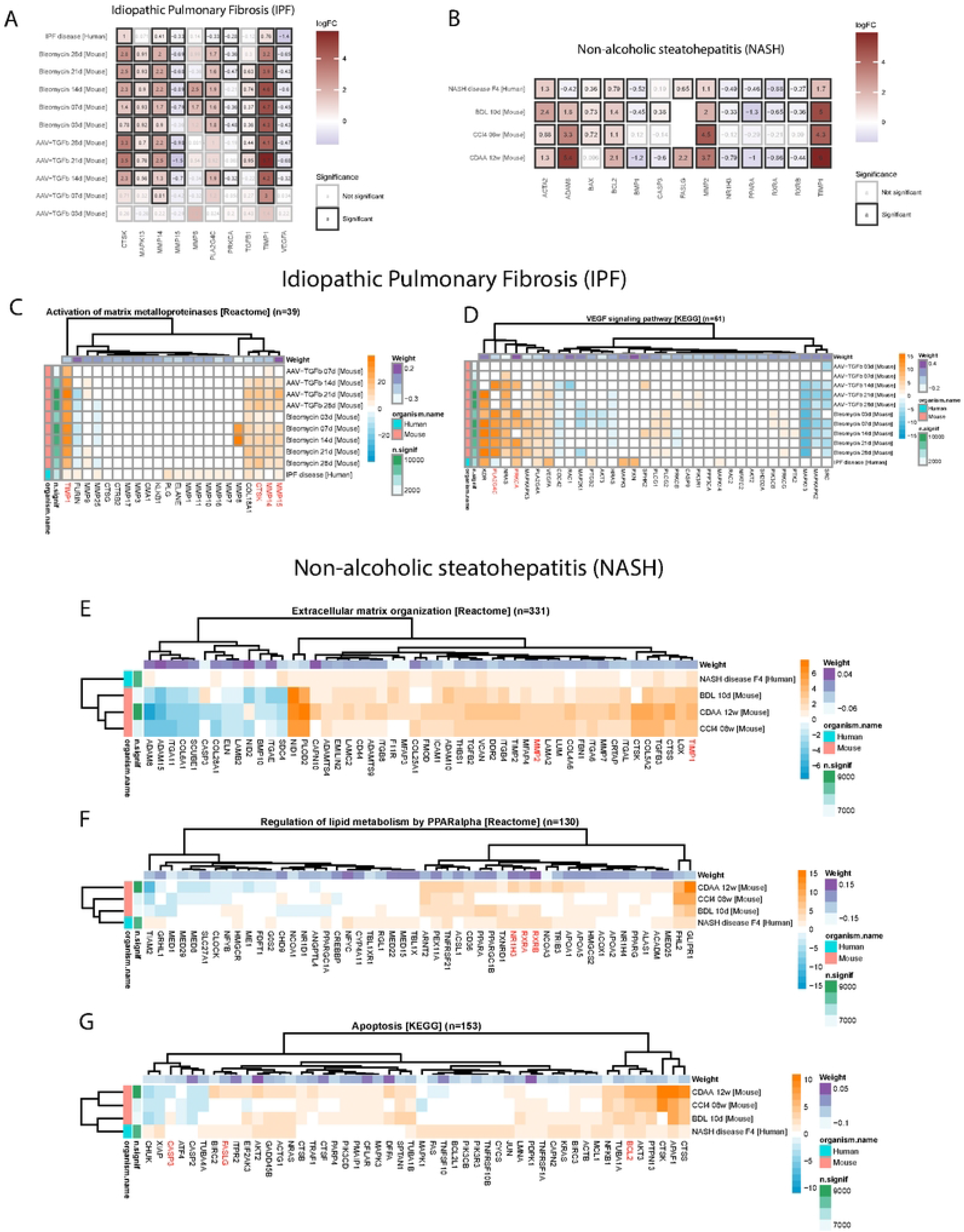
Assessment of gene contributions in hallmark pathways in human IPF and NASH using the IST gene heatmap: **(A)** Fold changes of disease-like states versus matched controls in logarithmic scale of a selection of relevant human genes and their murine one-to-one orthologs. Significance reported at 5% false discovery rate. **(B)** Analogous representation of fold changes for a selection of relevant genes in NASH. **(C)** Gene contribution heatmap obtained from IST, for the gene set “Activation of matrix metalloproteinases” as discussed in the IPF human disease context. Genes labelled in red are discussed in the main text. The heatmap scale represents gene contributions (%) for signature recapitulation. In orange, positive gene contributions imply that simulating the fold change of that gene helps bring human controls to IPF-like molecular profiles in that pathway, thus indicating agreement between species. In blue, negative gene contributions indicate disagreement, potentially implying opposite direction of change between humans and mice. In white, genes with low or no contribution; implies either no significant fold change, or low feature relevance in the context of classifying control versus human IPF in this pathway. The model weight scale describes the coefficient for each gene after fitting the linear predictor. Positive weights indicate genes that increase the disease score after upregulation, or equivalently, decrease the disease score after downregulation. Negative weights indicate genes that decrease the disease score after upregulation, or equivalently, increase the disease score after downregulation. **(D)** Gene contribution heatmap for the gene set “VEGF signaling pathway” in human IPF. **(E)** Gene contribution heatmap for the gene set “Extracellular matrix organization” in human NASH. **(F)** Gene contribution heatmap for the gene set “Regulation of lipid metabolism by PPARα” in human NASH. **(G)** Gene contribution heatmap for the gene set “Apoptosis” in human NASH.

### Gene-level comparison of the IPF disease models

We investigated the contribution of each individual gene in two IPF pathways that showed differences between the Bleomycin and the AAV-TGFβ1 model: Activation of matrix metalloproteinases pathway and VEGF signaling pathway (Figures 3C and 3D).

Within the activation of matrix metalloproteinases pathway, we observed strong upregulation of the fibrosis response marker *TIMP1* (Figure 3A). This upregulation was identified as highly relevant for the good recapitulation between human data and mouse models (Figure 3C). The upregulation of *Timp1* during a fibrogenic response is well established (Hall et al. 2003) and its Bleomycin-mediated as well as TGF-beta dependent activation has been shown (Strobel et al. 2015). These experimental data support the consistency between human and both mouse data sets observed by the IST analysis. Like *TIMP1*, the upregulation of metalloprotease *MMP14* and downregulation of *MMP15* (Figure 3A) showed alignment with human IPF gene expression changes across both mouse models (Figure 3C). IST highlighted the importance of *MMP8* upregulation (Figure 3A), which was specific to the Bleomycin model (Figure 3C). MMP8 has been already reported to be upregulated in both IPF patients and the Bleomycin model, and to correlate with the development of lung fibrosis, although its role in pathogenesis is not fully known (Pardo et al. 2016). In previous studies, Cathepsin K (CTSK), a member of the class of lysosomal-derived proteolytic enzymes, was found to be increased in fibrotic lung regions in patients and mice, and to provide a protective role by countering excessive deposition of collagen matrix in the diseased lung (Bühling et al. 2004). Indeed, IST provided evidence that the upregulation of *CTSK* gene expression (Figure 3A) is relevant for the alignment between human data and both animal models (Figure 3C).).

On the level of VEGF signaling, IST predicted that *VEGFA* is not the most influential gene (Figure 3D) to explain the differences in recapitulation of human IPF between the AAV-TGFβ1 and Bleomycin mouse models (Figure 2C). In fact, *VEGFA* expression was downregulated in humans and both mouse models (Figure 3A). Instead, IST results suggest that the difference between the mouse models in recapitulating human IPF gene expression was mostly explained by differences in regulation of *PLA2G4C* and *PRKCA* (Figure 3D). Indeed, we observed missing differential expression of *Pla2g4c* and *Prkca* in the AAV-TGFβ1 21d model, while they were up- and downregulated in the Bleomycin model, respectively (Figure 3A). PLA2G4C is part of the group 4 family members of phospholipidase A2 (PLA2) which is known as mediator of damaged-induced immune infiltration and vascularization. Cytosolic PLA2 is ubiquitously present in human lung and Pla2 knock-out mice had attenuated lung immune infiltration after Bleomycin treatment (Nagase et al. 2002). The good alignment in expression changes in *PLA2G4C* (Figure 3D), as well as its known role in vascularization, justified choosing the Bleomycin model over the AAV-TGFβ1 when investigating drug effects on VEGF signaling. On the other hand, the expression of the PKCα kinase had been previously shown to downregulate collagen expression via the MEK/ERK signaling pathway, together with findings of PKCα downregulation in fibrotic lung disease (Tourkina et al. 2005), which is consistent with IST’s prediction via *PRKCA*. As for potential disagreement between mouse and human, IST pinpointed that the upregulation of *Mapk13* in mice may require further investigation, as the same upregulation was not clearly found in the human reference.

### Gene-level comparison of the NASH disease models

In the previous section we found that IST predicts a high recapitulation of all animal models for the extracellular matrix organization pathway. This general agreement in IST was partly driven by several members of the pro-fibrotic tumor-derived growth factor beta 1 (TGFβ1) SMAD signaling pathway (Ghafoory et al. 2018), including a large contribution from the upregulation of tissue-inhibitor of metalloproteinases 1 (*TIMP1*) in humans and mice (Figure 3E). TIMP1 inhibits multiple matrix metalloproteinases (MMP), thereby preventing tissue remodeling and resolution of fibrosis (Iredale 2008). TIMP1 has also been described as a serum marker for advanced liver fibrosis in NASH patients (Yilmaz and Eren 2018) and is a known driver of fibrosis progression (K. Wang et al. 2013). Interestingly, Timp1-/- mice show increased liver fibrosis in CCl_4_-induced liver fibrosis (H. Wang et al. 2011), while in BDL fibrosis remains unaffected by the absence of TIMP1 (Thiele et al. 2017). IST did not predict this differential behavior because *Timp1* was upregulated in both CCl_4_ and BDL mice, as well as *TIMP1* in humans (Figure 3B). IST found agreement in the expression of Bone morphogenetic protein 1 (*BMP1*), see Supplementary File 1, due to its downregulation in humans and BDL and CDAA mice (Figure 3B). BMP1 processes multiple precursors of the extracellular matrix, as e.g., pro-collagen type I, and a Bmp1 splicing isoform has been shown to be a driver of disease progression in rat CCl_4_ models (Grgurevic et al. 2017). Since no *Bmp1* differential expression was found in CCl_4_ mice (Figure 3B), IST did not reproduce this claim in mice.

IST predicted good recapitulation for the Regulation of lipid metabolism by PPARα by the CDAA and BDL models, strongly influenced by the downregulation of *PPARA*, *RXRA*, *RXRB* and *NR1H3* (Figure 3F), as found in human NASH. These genes were however not differentially expressed in the CCl_4_ model (Figure 3B). PPARα can form a heterodimer with retinoid X receptors (RXRs) modulating gene expression of PPARα specific target genes via binding PPAR-response elements (PPRE). In the absence of PPAR ligands, the heterodimer acts as a co-repressor complex, while upon ligand binding, repressors are released and the PPAR-RXR heterodimer acts as a co-activator complex (Bougarne et al. 2018). Liver x receptor alpha (LXRα), encoded by the nuclear receptor subfamily 1, group H, member 3 gene (*NR1H3*) is another ligand-activated transcription factor of relevance in NASH, which controls lipid and glucose homeostasis (Voisin et al. 2020). There are and have been multiple drug discovery and clinical efforts to tackle MASH/MAFLD using small molecules targeting LXR receptors (Griffett and Burris 2023). LXRα phosphorylation has been shown to induce inflammation and fibrosis in the liver during high-fat diet feeding, while hepatic steatosis was found to be negatively regulated via LXRα (Becares et al. 2019). Due to the calculated importance of these genes in the molecular changes in human NASH, IST assigned a sensibly lower recapitulation to the CCl_4_ model in the PPAR pathway, where the metabolic NASH-driving component is lacking.

IST explained the varying degrees of recapitulation on the Apoptosis pathway in the animal models through noticeable contributions from known NASH biomarkers. While some markers showed overall strong positive recapitulation (*Bcl2*), others showed model-specific positive contributions: *Fasl* (CDAA), *Casp3* (BDL) and *Bax* (BDL, CCl_4_) (Figure 3G). FAS ligand (FASLG) induces apoptosis via binding to the FAS receptor and has been associated with NASH severity (Alkhouri et al. 2015). Accordingly, IST favored CDAA (Figure 3G) because it is the only mouse model with significant *Fasl* upregulation (Figure 3B). Cleaved caspase 3 is often used as a measurement of hepatocyte apoptosis in NASH (Feldstein et al. 2003). At the transcriptomics level, IST penalizes CDAA for having a significant *Casp3* downregulation, whereas it benefits BDL for showing upregulation (Figure 3B, 3G). The antiapoptotic regulator B-cell lymphoma 2 (BCL2) interacts and inhibits pro-apoptotic proteins, as well as it reduces apoptosis-related autophagy (K. Wang 2015). IST highlighted the importance of observing *Bcl2* upregulation in all the models, as found for *BCL2* in the human data (Figure 3B, 3G). Along these lines, Bcl2 inhibition has showed anti-fibrotic effects in mice (Teng et al. 2020), and BCL2 promotes resistance to pro-apoptotic stimuli in human hepatic stellate cells (Novo et al. 2006), underlining the key role of Bcl2 in liver fibrosis progression. Another element of the apoptotic cascade is the oligomerization of BCL2 associated X (BAX) and subsequent integration into the mitochondrial membrane, leading to membrane rupture and cytochrome c release, which triggers cleavage of pro-caspase 3 into active caspase 3 (Weiss et al. 2017). In line with this, IST found that the upregulation of *Bax* in the CCl_4_ and BDL models (Figure 3B) helped them recapitulate apoptosis as in human NASH (Supplementary File 1), since *BAX* was also upregulated in the human reference data.

### The In Silico Treatment framework includes features to assess and compare treatments for specific indications

In addition to assessing the quality of animal models to represent human disease through gene expression, IST can assess the molecular effect of treatment or recovery. To that end, IST uses the fold changes in gene expression between treated (or recovered) animal models versus those of the untreated animal model with disease. This reveals whether pathogenic gene expression changes are reverted by the treatment, and in addition may reveal potential unwanted effects of treatment.

### Fibrosis reversal following recovery from CCl_4_ induced liver fibrosis

To understand the capacity of IST to quantify the recovery of liver fibrosis we compared a dataset during the 4-, 8- and 12-week regression phase after an 8-week CCl_4_-induced liver fibrosis with the reversed human signatures, i.e. the fold changes between NASH fibrosis stages F4 and F0. Overall, recapitulation of the healthy states in the extracellular matrix organization pathway was 54% for 4-week and 68% for 12-week CCl_4_ recovery (Figure 4A), which is consistent with a partial, but not total, resolution of fibrotic phenotypes. While the gene product from the Acta2 gene, aSMA, as a measure of activated fibroblasts, rapidly decreased during recovery, the deposited collagen in the extracellular matrix was found to remain stable at high levels, even after 12 weeks of recovery (Supplementary Figure 3). In terms of gene contributions, our findings were analogous to those of NASH disease models: Downregulation of well-known regulators and components of the extracellular matrix like *Timp1* or *Mmp2* (Figure 4B) contributed to the positive recapitulation of human NASH (Figure 4C). Interestingly, IST identified some genes that disagreed in the reversal signatures from animal models at all three timepoints (Figure 4C). These included a disintegrin and metalloprotease 8 (*ADAM8*), downregulated in CCl_4_ recovery while upregulated in NASH fibrosis stage F4 to stage F0 reversal (Figure 4B). On one hand, *ADAM8* has been associated with chronic liver diseases, being increased in activated hepatic stellate cells, although the authors found no correlation with *MMP2* or *TIMP1*, and no changes in expression between fibrosis stages (Schwettmann et al. 2008). On the other hand, the neutralization of ADAM8 ameliorates acute CCl_4_-induced liver injury (S.-Q. Li et al. 2014).

**Figure 4.**
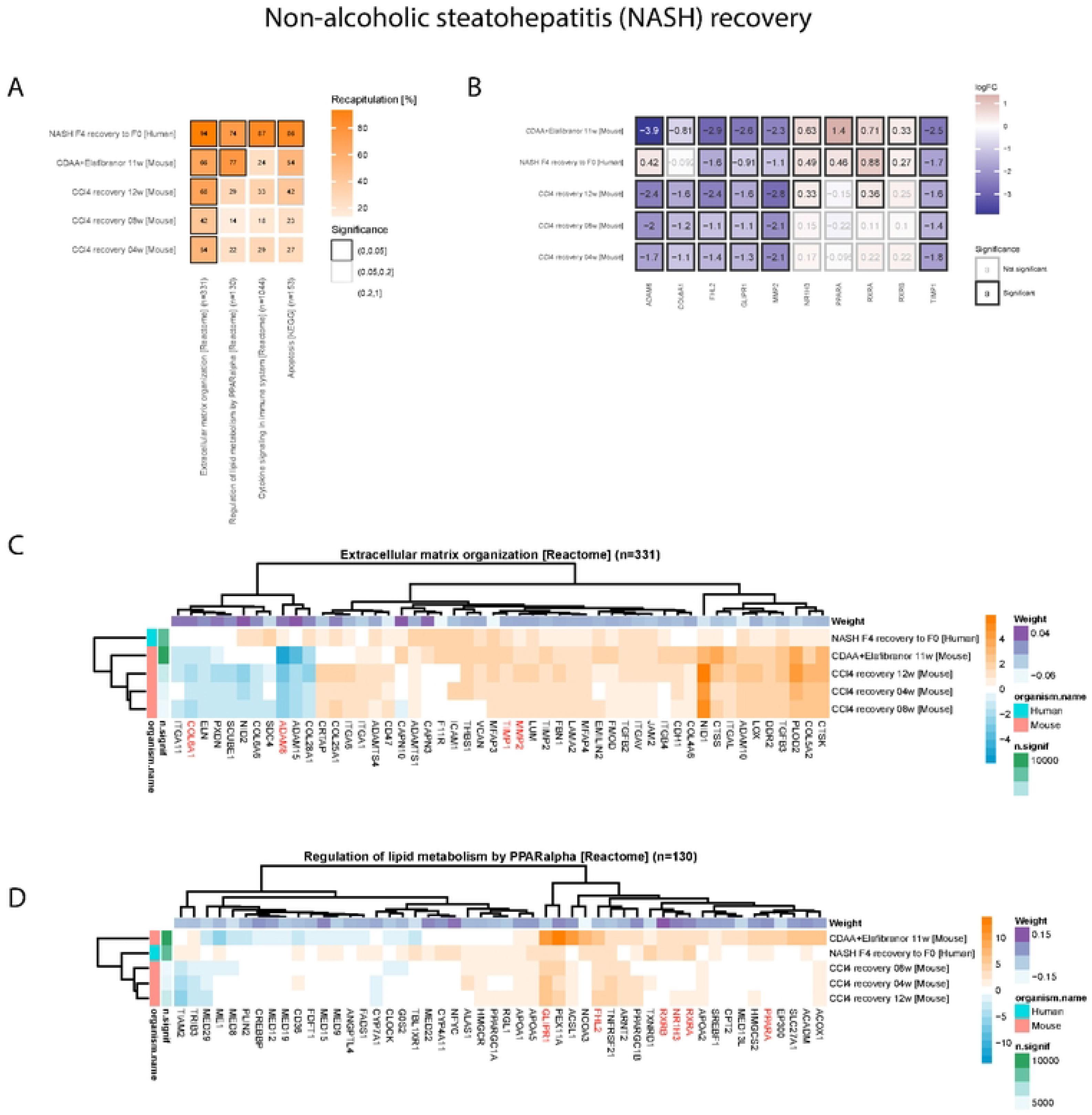
IST analysis to assess recovery from human NASH using the IST pathway and gene heatmaps: **(A)** IST pathway heatmap for the four human NASH hallmark pathways and the four NASH animal model signatures (three for recovery, one for treatment). We simulated fold changes on NASH F4 patients and expected good recovery signatures to bring the expression profiles closer to NASH F0 (ideal 100% recapitulation). **(B)** Fold changes of recovery versus disease-like states in logarithmic scale of a selection of relevant human genes and their murine one-to-one orthologs. Significance reported at 5% false discovery rate. **(C)** Gene contribution heatmap obtained from IST, for the gene set “Extracellular matrix organization”. Gene contributions (%) indicate whether changes in recovery or treatment models align with human NASH expression reversal. Positive (orange) contributions indicate changes in the same direction as the human reference data, whereas negative (blue) indicates changes in the opposite direction. **(D)** Gene contribution heatmap obtained from IST, for the gene set “Regulation of lipid metabolism by PPARα”.

### Elafibranor Treatment

One of IST’s potential applications is the *in silico* assessment of treatment effects. To that end we used Elafibranor, a PPAR agonist that has been considered as a potential treatment for NASH. Since we found CDAA was a robust model reflecting some of the main features of NASH (Figure 2D), we used Elafibranor as a NASH treatment in this model. Applying IST, we found that Elafibranor treatment showed strong recapitulations of healthy human expression patterns (Figure 5A) in liver fibrosis (66%), lipid metabolism regulation by PPARα (77%), apoptosis (54%), while moderate in cytokine signaling (24%). These findings are consistent with literature showing a strong effect of Elafibranor in an animal model of NASH and liver fibrosis (Hoek et al. 2021). Such recapitulations resembled those of 12-week CCl_4_ recovery, albeit PPARα regulation was sensibly lower in CCl_4_ recovery (29%).

For PPARα regulation of lipid metabolism, the high recapitulation in Elafibranor treatment on the CDAA mouse model (77%) even exceeds the recapitulation of CDAA itself as a disease model (62%,) or the recapitulation of the 12-week CCl_4_ recovery (29%) (Figure 2D). Genes of relevance for PPAR signaling that were identified by IST in NASH animal models (Figure 3F) *RXRA*, *RXRB* and *NR1H3* (LXR), show alignment between the human NASH reversal and CDAA mouse liver treated with Elafibranor (Figure 4D). We also found other genes contributing to Elafibranor’s positive recapitulation (Figure 4D): *PPARA*, upregulated in human NASH reversal and agonized by Elafibranor (Figure 4B), *GLIPR1*, downregulated in humans and mice (Figure 4B) and linked to stress-induced premature senescence as well as age-associated expression increase in mice hepatocytes (Doshida et al. 2023), and *FHL2*, downregulated in humans and mice (Figure 4B) and linked to hepatic fibrogenesis in humans and mice (Huss et al. 2013).

## Discussion

### Translational landscape

The fraction of drug candidates whose efficacy in animal models translated to clinical efficacy has remained steadily low in the last years, and it is unclear to which extent conclusions drawn from animal studies translate to human disease (Pound and Bracken 2014). Despite efforts in improving studies through better study design or bias control, translatability remains low due to biological differences and uncertainties between organisms. This remains an unsolved challenge amidst efforts to reduce unnecessary animal testing and improve animal welfare (Robinson et al. 2019). Thus, it is critical to leverage data and computational methods to aid the evaluation of suitable animal models for specific aspects of disease and avoid pitfalls in drug design.

Some computational tools can aid the process of animal model evaluation. Over-representation analysis of differentially expressed genes or rank-based gene set enrichment analysis (GSEA) represent essential tools to investigate gene expression changes in different species. However, while these analysis methods allow for the identification of affected pathways, they do not systematically integrate human and animal data. More sophisticated methods for the integration of human and animal model data have been developed. For example, the Found in Translation (FIT) method performs cross-species comparison using linear models (Normand et al. 2018). The Congruence Analysis for Model Organisms (CAMO) pipeline is another attempt, based on a Bayesian mixture model to quantify pathway-specific congruence scores (Zong et al. 2023).

### Methodological considerations

Here we present IST, a data integration tool to specifically address the quantification of key disease aspects in combination with a human reference. IST leverages transcriptomic readouts of humans and disease models to account for organismal similarities and differences. By design, IST provides information on the agreement between expression changes in human disease and animal models on a genomic, pathway, and gene level.

From a methodological perspective, IST relies on partial least squares models to define gene set-specific disease scores. This was a parsimonious model choice covering three key features. First, the outcome variable is computed via a linear predictor, which enables the explanation of changes in disease score through an exact decomposition in terms of individual gene contributions. Second, the number of features (transcripts) in a gene set can frequently exceed the number of samples used for model fitting, so a penalized method is required to handle the overdetermined system. The penalization was chosen not to induce sparsity, capturing subtle but coordinated changes in genes that may not reach univariate significance and letting the model coefficient assign an importance to that gene. Third, partial least squares provide natural choices for graphical sample representation and model diagnosis via its loadings and scores.

IST brings unique features on top of existing methods. A gene-wise predictive approach like FIT can help gain signal by finding new deregulated human genes starting from the mouse data, but does not quantify the degree of agreement per animal model off-the-shelf. The capability of computing a single number to represent pathway agreement already existed in CAMO, but there is no straightforward way to disentangle this measure by gene importance among the genes that agree or disagree. IST provides a single number per gene, integrating data on gene relevance for disease states classification within the pathway, change in mouse model and direction agreement. Another key feature is the quantification of the magnitude of change versus a desired outcome, which brings more nuance into the notion of agreement: changes can be too modest, just right or overly strong while always staying in the right direction. Furthermore, the formalism of IST also enables modelling quantitative outcomes in the human population, like disease stages or functional readouts.

### Informing decisions on animal model selection

We applied IST to compare the animal models for the selected pathways in IPF and NASH which complement the results of well-established gene set scoring methods such as the above mentioned GSEA. While GSEA assesses whether the genes in specific pathways show a statistically significant expression across conditions through the normalized enrichment score (NES), IST determines if gene expression changes in animal models align with those observed in the human reference via the percentage of recapitulation. Despite the GSEA and IST results layouts look similar, the rows displaying data on animal models in IST (Figure 2C, 2D) are already integrated with the human disease reference, whereas they are independent from the human reference in GSEA (Figure 2A, 2B).

### IPF study

We applied IST to assess 6 hallmark IPF features in the Bleomycin and AAV-TGFβ1 mouse models. IST captured the time-course component for optimal timepoint selection in a more insightful way than GSEA: IST predicted that earlier timepoints had lower recapitulation, and that both AAV-TGFβ1 and Bleomycin can recapitulate human molecular signatures of IPF in at least 4 out of the 6 selected pathways if the appropriate time point is selected. IST suggested that d14, d21 (Bleomycin) and d21 (AAV-TGFβ1) are sound timepoints in which both models recapitulate features of human IPF, with average recapitulations of 49.2%, 43.2% and 47.5% over the 6 pathways. The peak recapitulation in both models at d21 in extracellular matrix organization is in line with results published by the American Thoracic Society, which reported fibrosis appearing between days 14 and 28 after Bleomycin treatment (Jenkins et al. 2017).

The demonstrated clinical concept of Nintedanib treatment, together with the controversial role of VEFG signaling in IPF, provided a good opportunity to illustrate the value and granularity of IST. Some reports have linked increased, and potentially aberrant and overshooting neovascularization to increased Bleomycin-induced injury (Lee et al. 2008; Iyer et al. 2015). However, other authors argued that VEGF signaling after lung injury may act in an anti-fibrotic fashion, thereby being beneficial for prolonged survival and that lower expression of VEGF was correlated with a worse prognosis (Murray et al. 2017). Interestingly, the authors further demonstrated the antifibrotic role of VEGF in mice after Bleomycin treatment by attenuating collagen accumulation and lung remodeling. The IST results quantified that the Bleomycin induced injury in mice resembled VEGF-associated gene expression changes in human IPF more closely than the AAV-TGFβ1 mouse model. Based on our results, we speculate that TGFβ1-expression does not induce the same degree of vascular damage or injury-mediated re-vascularization as that observed upon Bleomycin-mediated lung injury. While this does not invalidate the AAV-TGFβ1 model, our *in silico* and *in vivo* treatment data support the hypothesis that parts of Nintedanib’s therapeutic effects on lung function might be more closely recapitulated in the Bleomycin model.

### NASH study

We applied IST to assess 4 hallmark NASH features in the CDAA, CCl_4_ and BDL mouse models. Overall, IST predicted CDAA as the best model to recapitulate the molecular signature of human fibrosis stage F4 for our selected group of 4 pathways, with an average recapitulation of 86%, followed by BDL (61.5%), and CCl_4_ (41.5%). CDAA also entailed the largest number of deregulated genes at the transcriptomics level (Supplementary Figure 1F). Fibrosis was the best recapitulated NASH aspects for the three models, which was expected given our focus on human fibrosis stage F4 versus F0. Our findings in apoptosis and cell death highlight the potential of computational tools like IST to strengthen standard scoring tools like the non-alcoholic fatty liver disease activity score (NAS) with apoptotic markers (Yanguas et al. 2016).

We showcased the capabilities of IST to assess treatments for human NASH. IST predicted the partial resolution of liver fibrosis in the CCl_4_ mouse model after 12 weeks of recovery, as a positive control. IST also recognized the strong anti-fibrotic effect of the PPAR agonist Elafibranor, as well as the risk of overshooting the PPARα activation. Elafibranor has recently been tested in a phase 3 clinical trial in patients with NASH and fibrosis, but failed to demonstrate a significant effect on NASH resolution as a monotherapy (GENFIT 2020). Taking everything together, IST added new evidence to the hypothesis that despite its strong anti-fibrotic effect, *PPARα* over-activation in animal models is among the plausible causes for Elafibranor’s lack of translation to the clinic (Rodriguez et al. 2018). This highlights the importance of integrating human and animal data for an early translatability assessment.

### Common findings between IPF and NASH in fibrotic disease

Since IPF and NASH fall under the common umbrella of fibrotic diseases, we expected to find commonalities from their analyses with IST. On one hand, *TIMP1* is a well-known fibrosis marker in both indications, for which IST quantified a substantial positive contribution, discussed in the context activation of matrix metalloproteinases and extracellular matrix organization. On the other hand, we discussed the role of *CTSK* in recapitulating the activation of matrix metalloproteinases in human IPF, but IST also underlined a positive contribution by *CTSK* in recapitulating extracellular matrix organization and apoptosis as they occur in human NASH. There is increasing evidence about the participation of cathepsins in liver disease pathophysiology and they are being investigated as biomarkers (Ruiz-Blázquez et al. 2021), and *Ctsk* has been found induced by the knockdown of the transcription factors *Elf3* or *Glis2* in mice in the context of hepatocyte reprogramming (Loft et al. 2021). Taken together, these findings suggest that CTSK may also play a role in human NASH and may deserve further examination.

### Assumptions and limitations

From the methodological perspective, the main assumptions behind IST when translating between species are: (i) the orthology mapping has enough coverage and quality to simulate enough changes on humans based on a one-to-one gene translatability, (ii) differential changes exist in both species, and (iii) the tissues are comparable in terms of cell composition. We checked to what degree such assumptions hold. Regarding point (i), on average, IPF and NASH animal model signatures had 21 873 and 14 826 transcripts, which mapped to 14 005 and 12 047 ortholog human genes, leading to a coverage of 64% and 81%. The IPF and NASH human references had 15 293 and 19 352 genes, out of which 12 425 (81%) and 12 570 (65%) had a mouse ortholog. The fact that we observed good overall recapitulations in the animal models, sometimes even exceeding the transcriptomics changes in humans, suggests that points (i), (ii) and (iii) were covered. The gene contribution heatmaps further support points (ii) and (iii) since contributions were mostly positive and in line with known disease markers.

IST heavily relies on the quality of the human reference data for model fitting, and specifically its data type, here bulk transcriptomics data for its broad availability. Thus, IST will only detect effects that are noticeable at that molecular level and resolution. The gene-level quantification was a valuable feature to detect specific instances in NASH where IST did not detect known regulation events. For instance, IST was unable to distinguish isoform-specific effects for *Bmp1*, for which paired end sequencing would be more adequate. IST did not find model-specific differences between CCl_4_ and BDL in TIMP1 regulation in the context of fibrosis, since *Timp1* was similarly upregulated in both models. IST could not account for Casp3 cleavage when evaluating the alignment between mouse and human apoptosis, and only evaluated *Casp3* deregulation at the transcriptomic level. IST highlighted potential disagreement between humans and mice in fibrosis resolution because conflicting changes in *ADAM8*, where changes in human disease may be clearer at a single cell resolution. These findings underline the importance of considering the trade-off between technological advantages and limitations behind the molecular data used for model selection.

### Conclusions

In summary, IST is a data integration computational approach that quantifies the alignment of changes in transcriptomic profiles in animal models and treatments to those of human disease. The roles of the animal and the human data are non-symmetric: IST is anchored on the human reference, where it learns the pathway-level differences in disease using the gene expression values, and only a signature of fold changes from animal or preclinical data is needed to simulate their effect in humans. IST was successfully applied to a smaller microarray dataset and a larger RNA-seq study, highlighting its robustness across platforms and sample sizes. IST is highly explainable since its decisions can be traced back to the gene level contributions. We found genes with key pathophysiological roles in humans and animals among genes with largest contributions. The rigorous data integration cannot be achieved using GSEA, where the effects of gene direction, effect size and significance are not combined off-the-shelf between both species. IST’s findings on two major indications, IPF and NASH, were supported by literature and by newly generated data, at the gene and pathway level. This showcased the potential of IST to make data-driven choices in the selection of the most appropriate animal models, hereby reducing costs and reducing ethical considerations in pre-clinical animal model research.

## Materials and Methods

### Human IPF reference expression data

Human IPF microarray data was obtained from the GEO (Gene Expression Omnibus) entry GSE47460 and subsampled according to the procedure specified in Wang and colleagues (Y. Wang et al. 2017). Raw microarray data was preprocessed by averaging the probe intensities for probes that represent the same gene, and further processed to obtain normalized gene expression levels.

Principal Component Analysis (PCA) on human expression data was performed using the pcaMethods R package version 1.78.0 (Stacklies et al. 2007). The following settings were applied: method = “nipals”, scale = “uv”, center = TRUE. Descriptive plots used the first and second principal components.

To attain class balance and focus on the common molecular features of the heterogeneous IPF landscape, IPF patients were subsampled to a representative selection by computing the medioids on the dimensionality-reduced principal components. The most representative IPF patients (medioids) were selected by compressing their expression profiles into *m* = 10 principal components (chosen *m* in 1, 2, …, 10 as the one maximizing the explained variance in prediction *Q*^2^ metric in a 5-fold cross-validation), computing all pairwise Euclidean distances between IPF patients, and picking the 12 IPF patients with the lowest average distance to the rest of patients. After balancing, limma v3.42.0 (Ritchie et al. 2015) was applied to calculate differential expression between control and IPF patients.

### Human NASH reference expression data

Human NASH RNA-sequencing data was obtained from the GEO entry GSE162694 (Pantano et al. 2021). Raw counts were preprocessed to obtain normalized gene expression levels. Differential expression was assessed between participants in fibrosis stages F4 and F0 using limma v3.42.0 on voom-normalised read counts. The NASH human recovery signature from F4 to F0 was obtained by flipping the sign of each fold change.

### IPF disease model data

#### Expression data

Two murine IPF preclinical models were evaluated in a single experiment: the Bleomycin and the AAV-TGFβ1 models, as published in the GEO entry GSE195773 (Strobel et al. 2022). After acclimating for one week, mice received intratracheal administration of either 2.5 × 10^11 vg of AAV-TGFβ1 or AAV-stuffer, 1 mg/kg Bleomycin, or NaCl solution in a volume of 50 µL. Mice were sacrificed at five timepoints: day 3, 7, 14, 21 and 28. Differential expression analysis was performed using Limma and the matrix of voom-normalized read counts (Ritchie et al. 2015). We compared each model versus its day-matched control by timepoint: day 3, 7, 14, 21 and 28. This led to 5 animal model signatures for the Bleomycin model and 5 signatures for the AAV-TGFβ1 model.

#### Lung capacity study

We performed a separate experiment to specifically assess the effect of Nintedanib in lung capacity on the Bleomycin and the AAV-TGFβ1 models, using C57BL/6JRj animals from Janvier. Mice were used in an age between 10-12 weeks. For both models, Bleomycin or TGFβ1 AAV (AAV6.2 (2.5E+11 VG/animal) were administered i.t. on day 0 and mice were sacrificed on day 21. Nintedanib was given 50mg/kg, p.o., b.i.d. Animal experiments were ethically approved by the Regierungspräsidium Tübingen, Germany; license: 16-028 and 18-032. Lung function was measured as described in (Weckerle et al. 2023).

#### NASH disease model data

Three murine NASH preclinical models were evaluated in four newly generated experiments.

#### Experimental design and RNA sequencing

The first experiment included a CDAA (choline-deficient, L-amino acid-defined) diet-based model in a cross-sectional study. It is expected that animals fed this diet develop pronounced liver steatosis and a certain degree of inflammation, with an addition of cholesterol to aggravate liver fibrosis. Janvier C57Bl/6JRj mice with an age of 8-9 weeks were fed with either choline-supplemented l-amino acid-defined (CSAA) Control E15668-04 or with CDAA 1% Cholesterol E15666-94 (https://www.ssniff.com) for 12 weeks. Animals were then sacrificed to extract and sequence RNA. 200ng of RNA were used with TrueSeq mRNA stranded Single Index protocol. Library was sequenced on HiSeq3000 with single end reads 85Bp reads + 7 index.

In a second experiment, the same CDAA model was used to test the experimental anti-fibrotic compound Elafibranor. Animals were treated with vehicle (0,5% Natrosol/0,015%TWEEN 80 in 5 mL/kg) or 15mg of Elafibranor (Genfit 505) bid from day 10 to the end of the experiment. Animals were sacrificed after 11 weeks with and without Elafibranor treatment under the CDAA diet. 250ng RNA was used as input for NEB mRNA_dual Index. Sequencing was performed on HiSeq4000 with 75bp single end + 8bp index.

The third experiment ran the CCl_4_ (carbon tetrachloride) liver toxicity model in a time-course design. Janvier C57Bl/6JRj mice with an age of 8-9 weeks were fed ad libitum with standard diet (KLIBA 3438). Control animals in the healthy group were fed with olive oil whereas animals in disease group were fed with 10ml/kg olive oil dilution of CCl_4_ with increasing dose: 0.875ml/kg at day 1, 1.75ml/kg during week 1-3, 2.5ml/kg during week 4-6 and 3.25ml/kg from week 7-10. A mouse subgroup was sacrificed after 8 weeks of CCl_4_ administration to obtain an animal model signature by comparing it to matched controls. Subsequent groups were left for 4, 8 and 12-week recovery to obtain three disease recovery signatures, comparing to the 8-week CCl_4_ group before recovery. 200ng of RNA were used with TrueSeq mRNA stranded Single Index protocol. Library was sequenced on HiSeq3000 with single end reads 85Bp reads + 7 index.

The fourth experiment performed bile duct ligation (BDL) or sham surgery in a time-course study. 70 male CD1 mice (8wks old at study inception) were purchased from Charles River Laboratories, US. Mice were acclimated under standard housing conditions on standard diet for 1wk prior to study initiation. The study was conducted in compliance with Boehringer Ingelheim IACUC protocols. All mice were administered a single dose of Buprenorphine HCL (0.1mg/lg) ≥60min prior to surgery. Mice were then anesthetized with a mixture of 2-3% Isoflurane + 1L/min oxygen. For BDL, the common bile duct was exposed through a midline abdominal incision, isolated from the surrounding tissue and occluded using two 5-0 sterile sutures placed 2-3 mm apart with the upper suture proximal to the hilum. The bile duct remained intact. Sham animals underwent identical surgical procedures whereby the tissue surrounding the bile duct was manipulated but without obstruction. The abdominal incision was closed, and mice regained consciousness quickly under post-operative supervision and returned to home cages for the duration of the study and maintained on standard rodent chow and water diet. Mice were monitored daily for health and euthanized per timepoint under isoflurane. Animals were sacrificed at 3, 5, 7, 10, and 14 days post surgery. Livers were collected and saved directly into RNA-later solution. Livers in RNA-later were kept at 4°C for 24hrs then transferred frozen at −80°C. Liver tissue was homogenized (Tissue Lyser II, Qiagen) using lysis buffer (TRIzol Reagent, Invitrogen). Total RNA was extracted from liver (PureLink RNA Mini Kit, Invitrogen), purified of gDNA (PureLink Genomic DNA Mini Kit, Invitrogen) and checked for quality and concentration (NanoDrop Eight Spectrophotometer, ThermoScientific). RNA quality analysis was performed using dilute purified RNA (GeneAMP PCR System 9700, Applied Biosystems) and (2200 TapeStation, Agilent Technologies). Samples with RNA Integrity Number less than 7.0 were not included in analysis. Samples were shipped to BGI Tech Solutions, (Hong Kong China) for next generation sequencing. Sequencing libraries were built according to the manufacturer’s procedures for the TruSeq polyA kit. Paired-end sequencing was performed on an Illumina HiSeq 3000 to a depth of roughly 25 million reads, with a read length of 100 bases.

### Data processing and differential expression

The pipeline for primary processing of NASH animal model RNA-Sequencing measurements has been previously described in detail (Söllner et al. 2017). We used the mouse reference genomes from Ensembl 84/GRCm38 (http://www.ensembl.org). Reads were mapped using the STAR aligner (Dobin et al. 2013). The gene expression was calculated using Cufflinks (Trapnell et al. 2013). Gene quantitation was performed with RSEM for generation of TPM and feature counts for generation of counts used in downstream analysis. Differential expression analysis was performed using Limma and the matrix of voom-normalized read counts (Ritchie et al. 2015).

Two kinds of signatures were obtained from differential expression contrasts: animal model signatures, when the contrast compared challenged animals to control animals, and treatment signatures, when the contrast compared challenged and treated animals versus challenged animals.

In the first CDAA study, we obtained one animal model signature of CDAA versus CSAA diet at 12 weeks. In the second CDAA study, we obtained one treatment signature from the CDAA diet with versus without Elafibranor treatment at 11 weeks. In the CCl_4_ study, we obtained one animal model signature comparing 8 weeks of CCl_4_ administration versus matched controls, and three treatment signatures comparing 4, 8 and 12-week recovery versus the 8- week CCl_4_ group. In the BDL study, we obtained one animal model signature by focusing on day 10 BDL versus sham surgery as the standard timepoint.

### Histological analysis in CCl_4_ study

To assess morphological changes in liver after the CCl_4_ challenge, a histological analysis was used to calculate values describing degree of fibrosis, steatosis, and the area with αSmooth Muscle Actin (αSMA) expression in histological images. Images were taken from paraffin sections of mouse liver, stained by a Masson trichrome method and an αSMA staining. Slides were systematically scanned with a Zeiss AxioScan.Z1 microscope (20x magnification) and exported with 1:2 scaling as images in TIF-format. In these images, the liver sections were segmented, and the area covered by liver then cut into mosaic tiles of size 1024 by 1024 pixels (from 160 to 716 tiles per slide). Shape information of the liver section for each tile was saved in images alpha channel for reuse during image analysis. Image analysis for all slides was done using HALO, a digital pathology software by Indica Labs (Corrales, NM, USA) that directly reads original czi-files. The Area Quantification Module was adapted to the αSMA and Masson staining and the whole tissue was analyzed. Total area with typical blue Masson staining was determined and used in the calculation of a value corresponding to Collagen-content. Total area with typical red RefineRed marker was determined and used in the calculation of a value corresponding to αSMA-content. The Vacuole Quantification Module was adapted to the Masson staining and used for the detection of vacuoles. Data were summarized with Tibco Spotfire, analysis was done with GraphPad Prism. The color deconvolution could not sufficiently separate the aSMA marker (stain 1) and the blue counter stain (stain 2). Therefore, the area with aSMA was corrected by subtracting double stained areas. This was done in Spotfire, calculating [% Stain 1 Positive Tissue] - [% Colocalized Tissue (stain 1 and 2)].

### Gene annotations and mappings

#### Orthology mapping and primary gene identifiers

One-to-one orthologs were retrieved from the ENSEMBL (Yates et al. 2019) homology resource (jan2020.archive.ensembl.org) between Homo sapiens and Mus musculus ENSEMBL identifiers were used as primary throughout the analysis. Entrez gene symbols were mapped to ENSEMBL using biomaRt 2.42.0, archive version sep2019.archive.ensembl.org (Durinck et al. 2009).

#### Gene set and pathway data

Pathway-related gene sets were obtained from KEGG Release 96.0+/11-20 (Kanehisa et al. 2022). The selection of Reactome pathways (Gillespie et al. 2021) came from MSigDB version 7.0, C2 category, “CP:REACTOME” subcategory (Liberzon et al. 2015).

#### Gene set enrichment analysis

Gene Set Enrichment Analysis, or GSEA (Subramanian et al. 2005), was performed via the GSEA() function from the clusterProfiler R package version 3.14.2 (Yu et al. 2012), using pathway related gene sets mentioned above. For this analysis, genes were ranked by their fold changes. Mouse genes from animal model data were previously mapped to its human orthologue as described above. We excluded gene sets smaller than 15 genes from our analysis, while no upper limit on size was set. For each ranked list, the following parameters were used: by = “fgsea”, exponent = 1, pAdjustMethod = “BH”, nPerm = 100000, seed = TRUE.

### In Silico Treatment

#### Input data

IST requires the following input data: molecular readouts for the human disease, fold changes for the animal models, an orthology mapping and a list of gene sets of interest. Their respective indexing notation is described in Table 1: human samples are denoted by *i* (ranging from *i*_1_ to 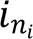), human genes by *j* (*j*_1_ to 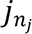), the quantitative values of disease scores by *k* (*k*_1_ to 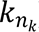), gene sets by *s* (*s*_1_ to 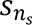), and statistical contrasts by *t* (*t*_1_ to 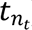). The variables mentioned throughout the methods that build on this notation are summarized in Table 2.

**Table 1.** Indexing notation for the human, animal, orthology and gene set (pathway) data.

| Entity | Index | First element | Last element |
| --- | --- | --- | --- |
| Human sample | $i$ | $i_1$ | $i_{n_i}$ |
| Human gene | $j$ | $j_1$ | $j_{n_j}$ |
| Disease score value | $k$ | $k_1$ | $k_{n_k}$ |
| Human gene set | $s$ | $s_1$ | $s_{n_s}$ |
| Contrast in animal models | $t$ | $t_1$ | $t_{n_t}$ |

**Table 2.** Description of variables as used in the In silico Treatment models.

| Variable | Description |
| --- | --- |
| $x_{ij}$ | Log2 expression value of the $j$-th gene for the $i$-th sample |
| $y_i$ | Disease score of the $i$-th sample |
| $g_k$ | Set of samples with a disease score of $k$ |
| $r_{jt}$ | Log2 fold change of the $j$-th (ortholog) gene in the $t$-th disease model signature (zero for non-significant genes) |
| $\hat{y}_{is}$ | Predicted disease score for the $i$-th sample in the $s$-th gene set regression model |
| $\beta_{js}$ | Coefficient of the $j$-th gene in the $s$-th gene set regression model |
| $\varepsilon_{is}$ | Error in the $i$-th sample within the $s$-th gene set regression model |
| $x'_{ijt}$ | Log2 expression value of the $j$-th gene for the $i$-th sample after in silico treatment with the $t$-th signature |
| $\hat{y}'_{its}$ | Predicted disease score for the $i$-th sample in the $s$-th gene set regression model after in silico treatment with the $t$-th signature |
| $\delta_{jts}$ | Change in prediction within the $s$-th gene set regression model associated to the $j$-th gene in the $t$-th signature |
| $\Delta_{ts}$ | Change in prediction within the $s$-th gene set regression model associated to the whole $t$-th signature |
| $\Delta_{0s}$ | Ideal change in prediction (recapitulation) of the $s$-th gene set regression model |
| $\delta f_{jts}$ | Percentage of ideal recapitulation within the $s$-th gene set regression model associated to the $j$-th gene in the $t$-th signature |
| $\Delta f_{ts}$ | Percentage of ideal recapitulation within the $s$-th gene set regression model associated to the $t$-th signature |

#### Predictive modelling of human data

The quantitative nature of IST relies on regression models, able to predict the disease stage of arbitrary humane gene expression profiles. To fit predictive models, features (human gene expression readouts) *x*_*ij*_ were provided in a scale suitable for addition, such as log2- transformed expression values), with no missing entries or constant genes. We further defined the response variable *y*_*i*_, indicating disease stage. Based on disease stage, samples were stratified into sample groups *g*_*k*_. If only control and disease samples were available, we set *y*_*i*_ = ―1 for disease and *y*_*i*_ = 1 for controls, and defined two sample groups *g*_―1_ = {*i* | *y*_*i*_ = ―1}, *g*_1_ = {*i* | *y*_*i*_ = 1} accordingly (see notation in Table 2).

Partial least squares, or PLS (Mevik and Wehrens 2007) models were fit using the caret R package version 6.0-85, within each gene set separately, yielding a total of *n*_*s*_ models. Let *s* be a gene set with *l* genes, noted as *j*_1_,…,*j*_*l*_ without loss of generality. The disease scores *y*_*is*_ were expressed as:

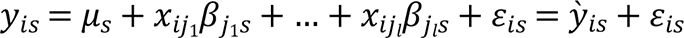

where *ỳ*_*is*_ is the predicted disease score for the *i*-th sample in the *s*-th gene set. The model coefficients 𝜇_*s*_ and 𝛽_*js*_ were fitted using method = “kernelpls”. Features were centered and unit scaled. For notation convenience, 𝜇_*s*_ includes all the feature centering and 𝛽_*js*_ includes the scale, i.e. is determined by dividing the model coefficient by the scaling factor of *x*_*ij*_. The number of components was selected from 𝐾 ∈ {1, 2, 3, 4, 5} using 5-fold cross-validation, repeated 20 times. Selection criteria was the minimum root mean squared error in prediction. The final model was fitted with the optimal 𝐾.

#### Fold change projection

A main step in IST is the projection of disease model signatures (fold changes associated with a statistical contrast *t*) onto human expression data. As detailed above, log2 fold changes were calculated following the limma convention of linear modelling (Ritchie et al. 2015). For each signature, only significantly deregulated genes with ∣log_2_ 𝐹𝐶∣ > 0.25 and false discovery rate 𝐹𝐷𝑅 < 5% (Benjamini and Hochberg 1995) were considered. Gene identifiers were mapped to one-to-one human orthologs, thus avoiding collisions of several animal genes mapping to the same human gene. Finally, the log2 fold change of an animal gene *j* with a human ortholog *j* within the *t*-th signature was denoted *r*_*jt*_, where *r*_*jt*_ = 0 if *j* was not significant in *t*. The projection of fold changes, which we refer to as *fold change simulation* or overlay, was then defined as follows (Table 2):

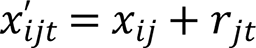

Two types of signatures were considered: disease models and treatments. Disease models compare challenged versus control animals, whereas treatment signatures compare treated challenged animals with untreated challenged animals. The choice of simulated human samples and reference samples was determined by the corresponding sample groups. When assessing disease models, the aim is to simulate the challenge from animals onto human samples in *g*_1_ and compare the outcome to those in *g*_―1_. The roles of *g*_―1_ and *g*_1_ are switched when assessing treatments. As a positive control for disease models, we included signatures obtained from the human reference data.

### Quantification of disease recapitulation

Here we define recapitulation as the similarity between samples with simulated fold changes and reference samples. Recapitulation was quantified by predicting the disease scores of simulated samples using the previously fitted PLS models. The ideal recapitulation in animal models within the *s*-th gene set (Table 2) was defined as:

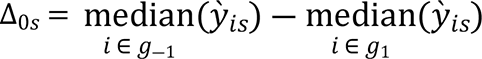

On the other hand, for treatments:

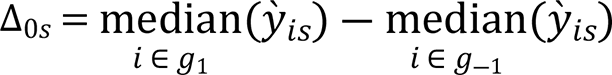

When simulating the fold changes onto the human samples in *g*_―1_ (animal models) or *g*_1_ (treatments), the predicted disease score change Δ_*ts*_: = *y*’_*its*_ ― *ỳ*_*is*_ is independent of *i*, as shown:

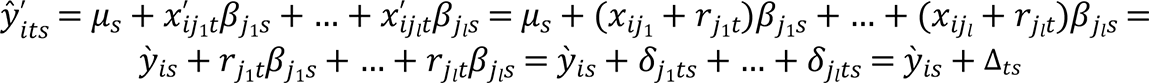

Therefore, the change can be expressed down to the gene-level contributions, defining *δ*_*jts*_ : = *r*_*jt*_𝛽_*js*_, which do not depend on *i*:

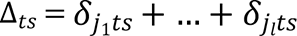

To give a reference on the magnitude of the gene contributions *δ*_*jts*_ and the whole signature changes Δ*ts*as a fraction of the ideal recapitulation, the following relative percentages were defined.

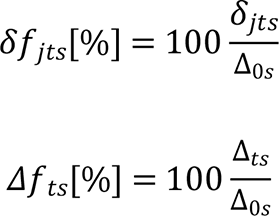

Those were easier to interpret and still verify that the overall recapitulation can be expressed as the sum of each gene’s contribution, i.e. 𝛥*f*_*ts*_[%] = *δf*_*j*1*ts*_[%] +… + *δf*_*jlts*_[%]. A recapitulation of 𝛥*f*_*ts*_ ≈ 100% would imply that the median disease scores of samples simulated with fold changes from signature *t* corresponds to that of the reference samples. Accordingly, gene-level contributions *δf*_*jts*_ further show which genes had more influence in the final recapitulation. This justified why IST predicted strong or weak recapitulations. Genes meeting two conditions would provide large contributions in the right direction (*δf*_*jts*_ ≫ 0, i.e. agreement): having a large, significant fold change in the disease model, and finding the same direction of change in the PLS model in human data. Conversely, genes with large contributions in the opposite direction (*δf*_*jts*_ ≪ 0, i.e. disagreement) would arise from strong changes in the disease model and the human data, but with opposite directions. Finally, genes would show little contribution (*δf*_*jts*_ ≈ 0) if either they were not differential in the disease model, or the PLS model found barely any changes in the human reference, or both.

To evaluate the statistical significance of recapitulation 𝛥*fts* of a signature *t* within a gene set *s* we devised a null model for size-matched signatures and computed their recapitulation. In each null trial, carried out per animal study, the identities of all the genes were shuffled, so that the original number of differential genes and their fold change distribution were preserved. If time points were present, this also kept longitudinal gene co-expression patterns. The empirical p-values (North, Curtis, and Sham 2003) for the observed 𝛥*fts* was then computed as 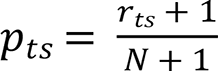, where *r*_*ts*_ was the number of null trials, out of 𝑁 = 1000, with a recapitulation as extreme as 𝛥*fts*. Empirical p-values were then adjusted for false discovery rate.

### Graphical representations

The predicted disease scores for untreated samples *ỳ*_*is*_ and their simulated counterparts *ŷ*^′^_*its*_ (Table 2) could be represented through gene set-wise boxplots. Keeping *s* fixed, *ỳ*_*is*_ were grouped in boxes by *g*_*k*_ and *ỳ*_*its*_ʹ by the signatures *t*. Every data point in the boxes corresponded to a sample *i*. The untreated samples would illustrate the reference ranges of disease scores for normal and disease states.

The overall gene set recapitulations 𝛥*f*_*ts*_ were represented in heatmaps using the pheatmap R package version 1.0.12, where the rows were indexed by the signature *t* and the columns by the gene set *s*. The signature with the original human fold changes would serve as a reference recapitulation. Optionally, we displayed hierarchical clustering of the rows and columns used Euclidean distances and the “complete” method in hclust(), to unravel patterns of similar and dissimilar recapitulations in gene and signature clusters (Everitt et al. 2014).

For each gene set *s*, a heatmap was drawn to depict the gene level contributions. Fixing *s*, the *δ*_*jts*_ values were arranged, indexing the rows by the signature *t* and the columns by the gene *j*. Again, the human signature would serve as a reference. Due to the large size of individual gene sets, only the top 50 contributing genes were displayed, defined as those with the largest sum 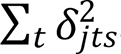. Optionally, hierarchical clustering was applied to highlight similar patterns in both gene and signature recapitulations.

## Acknowledgements

The authors thank David Lamb for useful early discussions and ideas on the topic. The authors thank Angela Lopez-del Rio for useful comments on the manuscript. The authors thank Glenn Gibson for his assistance with BDL surgical and study procedures. The authors thank Dagmar Knebel-Haas, Werner Rust, David Kind and Eleonora M. Capitolo for NGS data generation. The authors thank Stefano Patassini for his support throughout the publication process. The authors thank Piotr Radkowski for his support building interactive visualizations.

## Figure Legends

**Supplementary Figure 1.**
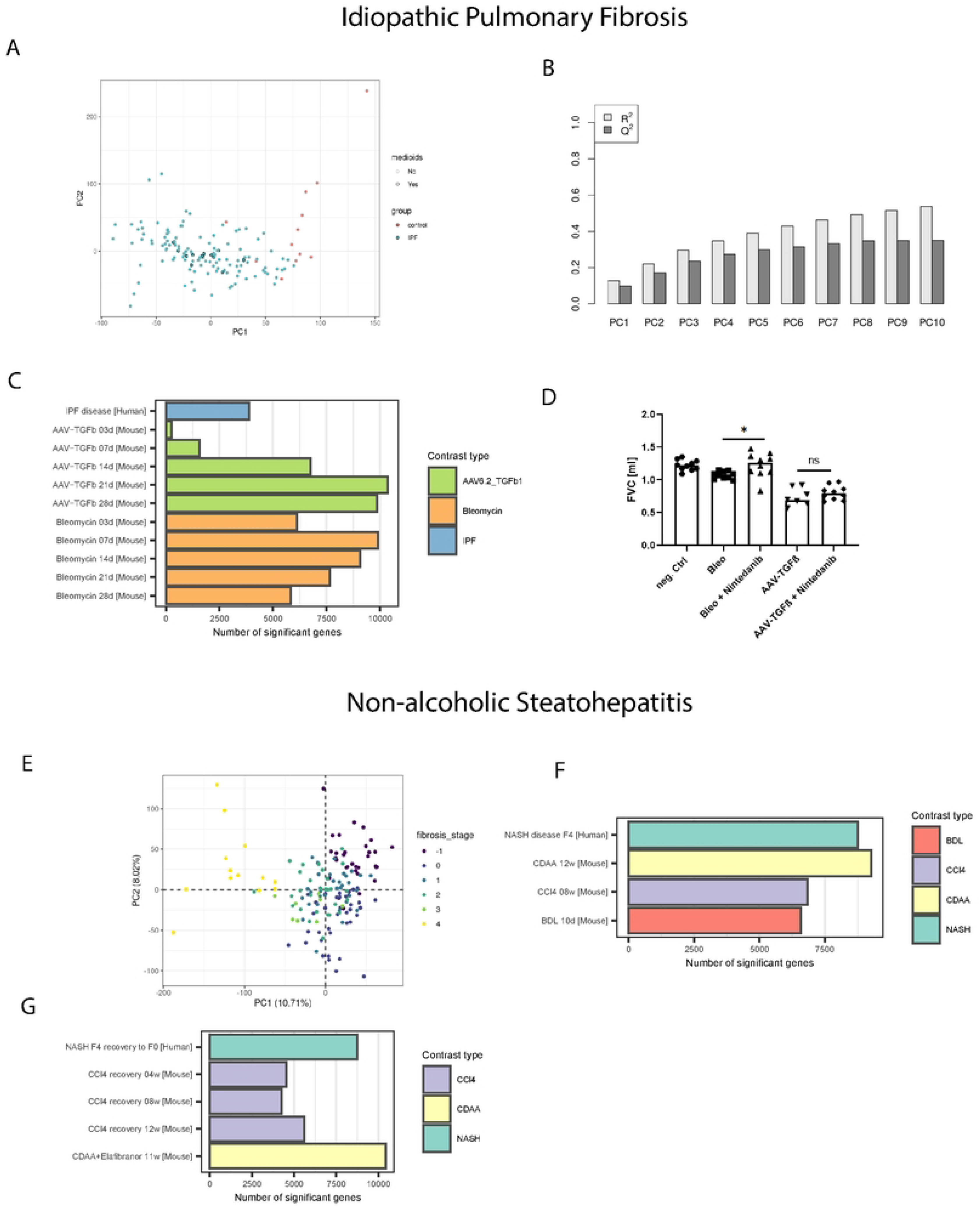
Descriptive statistics of the animal models for IPF and NASH, and human IPF and NASH data: **(A)** Principal components 1 and 2 for the IPF human reference data. IPF samples are shown in blue, while control samples are shown in red. Inclusion of individual IPF samples in the medoid subset is indicated by a black outline. **(B)** For principal components 1 to 10 in the IPF human reference data, cumulative percentage of variance explained (R^2^) and its cross-validated version (Q^2^). **(C)** Number of significant genes (absolute log fold change above 0.25 and false discovery rate below 5%) for the human IPF and the animal model signatures. **(D)** Lung forced vital capacity in the AAV-TGFβ1 and the Bleomycin models with and without Nintedanib treatment. **(E)** Principal components 1 and 2 for the NASH human reference data. The color scale represents the fibrosis stages. **(F)** Number of significant genes for the human NASH disease and the animal models. **(G)** Number of significant genes for the human NASH reversal and the animal recovery signatures.

**Supplementary Figure 2.**
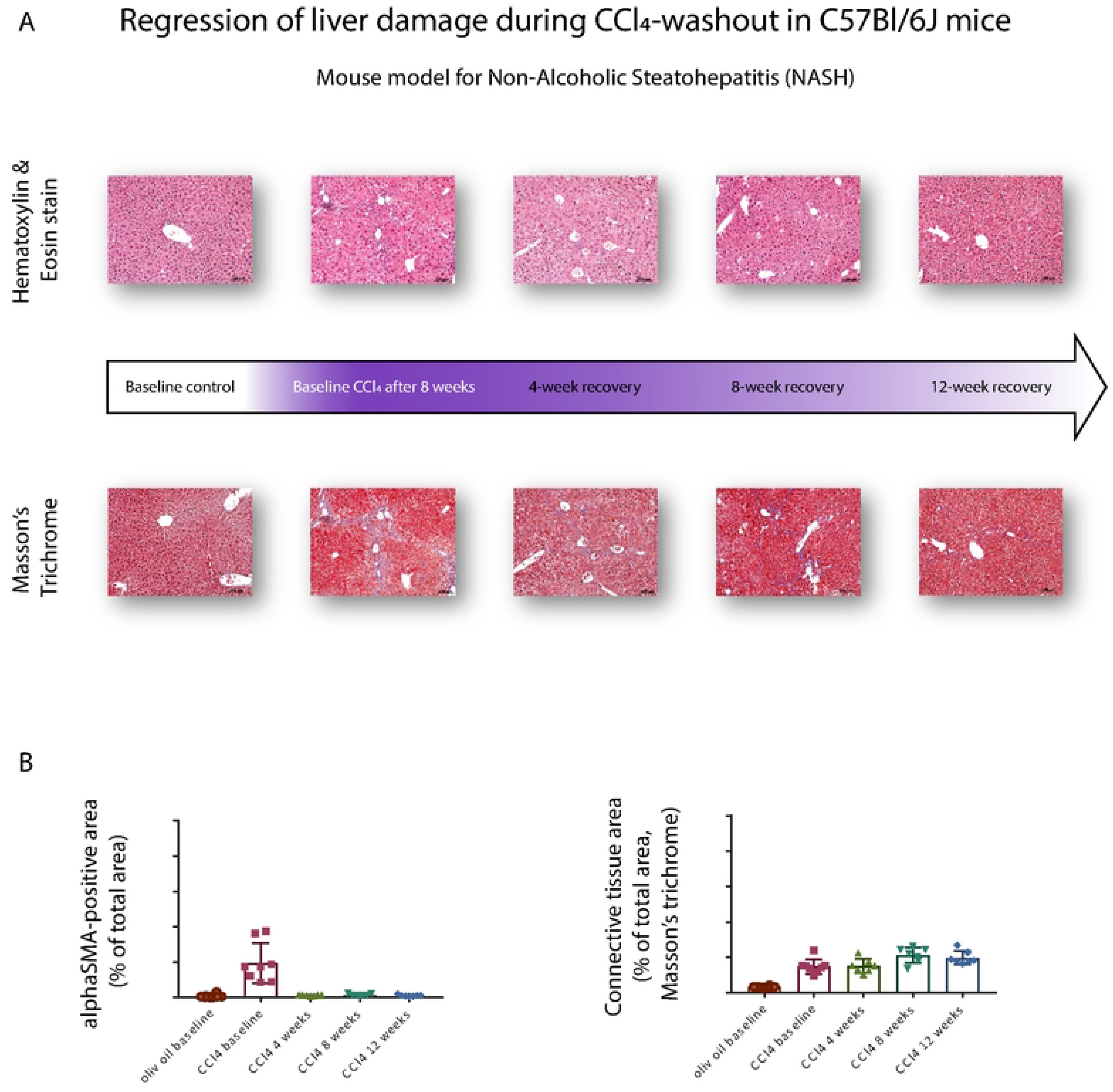
Regression of liver damage during CCl_4_-washout in mice. **(A)** Histological images of mouse liver during the baseline control, CCl_4_ challenge and subsequent 4, 8 and 12-week recovery (hematoxylin and eosin stain, Masson’s Trichrome). **(B)** aSMA and collagen area as computed from image data.

**Supplementary File 1 – Full gene heatmaps as computed with IST in the IPF and NASH use cases**. Zip file where all the genes belonging to each gene set were displayed, as opposed to the figures in the main text, which are limited to the top 50 genes. Plots include the evaluation of animal models in IPF and NASH, and the evaluation of treatments or recovery in NASH.

## Data and code availability

All the newly generated mouse sequencing data will deposited in GEO.

The code implementing the computational methods in IST is available as an R package called IST at https://github.com/bi-compbio/IST, with a vignette that describes the approach, implementation, and usage. IST also bundles an interactive R shiny app, available at https://github.com/bi-compbio/IST_browser, that displays an IST results object to prioritize signatures and pathways by recapitulation, and to compare signatures within pathways. The code and data to reproduce the results of this manuscript can be found at https://github.com/bi-compbio/IST_results

